# Optogenetic Modeling of Wake-Like Transcriptional Progression in Human SH-SY5Y Neuronal Cells

**DOI:** 10.64898/2026.02.27.708551

**Authors:** Shintaro Yamazaki, William Gee, Utham K. Valekunja, Akhilesh B. Reddy

**Affiliations:** Department of Systems Pharmacology & Translational Therapeutics, Perelman School of Medicine, University of Pennsylvania, Philadelphia, PA 19104, USA; Institute for Translational Medicine and Therapeutics, Perelman School of Medicine, University of Pennsylvania, Philadelphia, PA 19104, USA; Chronobiology and Sleep Institute (CSI), Perelman School of Medicine, University of Pennsylvania, Philadelphia, PA 19104, USA; University of Cambridge Metabolic Research Laboratories, Wellcome Trust-MRC Institute of Metabolic Science, Addenbrooke’s Hospital, Cambridge CB2 0QQ, UK; National Institute of Animal Biotechnology (NIAB), Opp. Journalist Colony, Near Gowlidoddi, Extended Q City Road, Gachibowli, Hyderabad - 500 032, Telangana, India

**Author notes:** Equal contributions (Shintaro Yamazaki, William Gee).

## Abstract

Sleep deprivation induces extensive transcriptional remodeling in the brain, yet it remains unclear how much of this response arises directly from sustained neuronal excitation versus systemic and behavioral confounds. Here we establish a reductionist human neuronal cell model to isolate the cell-autonomous consequences of prolonged depolarization. Controlled optogenetic stimulation of human SH-SY5Y neuronal cells elicited scalable calcium influx and robust induction of canonical activity-dependent genes, enabling precise manipulation of excitation history independent of receptor-specific pleiotropy. Time-resolved RNA sequencing revealed that sustained excitation does not produce a uniform transcriptional shift. Instead, responses partitioned into a small set of reproducible temporal modules including early-, mid-, and late-peaking induction, sustained induction, biphasic responses, and suppression, each associated with distinct functional and regulatory signatures. At the systems level, pathway activities evolved through three sequential transcriptional states, each defined by a distinct mixture of these temporal gene modules within individual programs. State transitions were directional and asymmetric across stimulation and recovery, consistent with cumulative excitation-history–dependent reorganization rather than simple reversal after stimulus offset. Cross-dataset mapping with *in vivo* mouse sleep datasets showed selective concordance between defined temporal modules and independent sleep-deprivation signatures. Together, these findings show that sustained neuronal excitation alone is sufficient to generate staged, wake-like transcriptional progression, linking gene-level temporal structure to higher-order regulatory states shaped by excitation history.

## Introduction

Sleep deprivation profoundly alters neuronal physiology and gene expression, reshaping transcriptional, metabolic, and proteostatic programs in the brain. Genome-wide studies consistently demonstrate the induction of immediate-early genes, activation of stress-responsive pathways, and metabolic reconfiguration following prolonged wakefulness ^1–3^. Sustained wake has been linked to proteostatic strain and the engagement of heat shock and unfolded protein responses ^4–6^. Although these molecular signatures are well characterized *in vivo*, attributing them specifically to sustained neuronal activity and elevated synaptic drive, core cellular correlates of wakefulness, remains challenging. In intact organisms, prolonged wake is accompanied by coordinated alterations in brain and body temperature, metabolic flux, glucocorticoid signaling, and inflammatory tone ^7–11^, potentially confounding the mechanistic interpretation of excitation-driven transcriptional remodeling. This is compounded by inverted sleep-wake profiles of nocturnal rodent models of sleep (relative to the solar cycle) compared to diurnal organisms including humans. Thus, while *in vivo* sleep-deprivation models establish organismal relevance, they limit mechanistic isolation of excitation-driven gene regulation.

Reductionist approaches have partially addressed this limitation. Primary neuronal cultures and cortical explants retain core molecular correlates of sleep-wake regulation, indicating that key transcriptional programs can arise within isolated neuronal assemblies ^12^. However, existing *in vitro* paradigms rely on pharmacological depolarization or neurotransmitter cocktails, which engage pleiotropic receptor signaling and offer limited control over excitation history ^12^. As a result, they cannot readily distinguish rapid activity-coupled induction from slower, cumulative remodeling that depends on prior excitation exposure.

Optogenetics provides temporally precise and reversible control of depolarization ^13–15^, enabling excitation itself to be treated as an explicit experimental variable while minimizing receptor-mediated confounds. SH-SY5Y cells, a human neuroblastoma-derived neuronal model, exhibit calcium signaling and activity-dependent gene induction upon depolarization and are widely used as a tractable human neuronal model ^16–18^. Their genetic tractability and compatibility with stable opsin expression provide a controlled platform for isolating excitation-driven transcriptional programs in a human context.

Here, we establish an optogenetically-controlled human neuronal model of sustained excitation and characterize its transcriptional dynamics across stimulation and recovery. Rather than eliciting a uniform transcriptional response, prolonged excitation organizes gene expression hierarchically: reproducible temporal gene modules assemble within functional programs, and their shifting mixtures define discrete, sequential transcriptional patterns over time. By isolating excitation history from systemic and receptor-level interference, this framework captures staged, wake-like transcriptional progression in a tractable human system and provides a mechanistic platform for dissecting excitation-history–dependent gene regulation relevant to sleep deprivation.

## Methods

### Cell Culture and Maintenance

SH-SY5Y human neuroblastoma cells were maintained in a 1:1 mixture of Opti-MEM (Gibco, 31985-047) and Ham’s F12 (Sigma, N6658) supplemented with 12% fetal bovine serum (HyClone, SV30180.03), 1% GlutaMAX (Gibco, 35050-038), 1% non-essential amino acids (Sigma, M7145), 1% penicillin–streptomycin (Sigma, P0781), and 0.2% MycoZap Plus (Lonza, VZA-2022). Cells were cultured in Corning T75 flasks at 37°C in a humidified incubator with 5% CO₂. Cells were passaged at 70–90% confluence. Culture medium was aspirated, cells were rinsed with pre-warmed phosphate-buffered saline (PBS), and detached using 3 mL trypsin–EDTA (Sigma, T3924). After detachment, trypsin was neutralized with complete medium, and cells were either reseeded for expansion or plated for transfection and downstream assays.

### Generation of Plasmids for Stable Expression of Opsins

Plasmids encoding opsin-based excitation constructs were generated by Gibson Assembly using a modified pEGFP-N1 backbone (Clontech) in which the GFP coding sequence was removed to accommodate custom inserts. The backbone contains a cytomegalovirus (CMV) promoter, pBR322 origin, neomycin/kanamycin resistance cassette, and SV40 origin. Coding sequences were amplified using Phusion Hot Start II DNA polymerase (Thermo Fisher Scientific) and assembled into the linearized backbone using NEB Gibson Assembly Master Mix (NEB, E2611S) according to the manufacturer’s instructions. Primer sequences used for amplification are provided in Supplementary Table S1. Two initial constructs were generated: CRIP, encoding Channelrhodopsin-2 (C128T mutant; ChR2(C128T)) fused to RCaMP1h (red genetically encoded calcium indicator 1h) followed by an internal ribosome entry site–puromycin resistance cassette (IRES-PuroR), and CHALIP, encoding ChR2(C128T) linked via a 2A self-cleaving peptide to enhanced Natronomonas pharaonis halorhodopsin 2.0 (eNpHR2.0), followed by IRES-PuroR. The ChR2(C128T) sequence was amplified from Addgene plasmid #20295 (deposited by K. Deisseroth). RCaMP1h was amplified from Addgene plasmid #42874 (L. Looger), and the IRES-PuroR cassette was amplified from Addgene plasmid #30205 (D. Kotton). The 2A-eNpHR2.0 construct was assembled using the N-terminal fragment from Addgene plasmid #22047 (E. Boyden) and the C-terminal fragment from Addgene plasmid #26966 (K. Deisseroth). The plasmid backbone was generated by restriction digestion of Addgene plasmid #22047 with EcoRI, KpnI, and EcoRV followed by gel purification. To compare excitation efficiency across opsins, the ChR2(C128T) sequence in CRIP was replaced with alternative channelrhodopsins, including Chronos (Addgene #62726, E. Boyden) and CoChR (*Chloromonas oogama* channelrhodopsin; Addgene #59070, E. Boyden). Each opsin was fused to the RCaMP1h-IRES-PuroR cassette amplified from CRIP and assembled into the same pEGFP-N1-derived backbone using Gibson cloning. Assembly reactions were transformed into chemically competent E. coli (Stbl3 or NEB C2527I), and transformants were selected on LB agar containing 50 µg/mL kanamycin. Positive clones were screened by colony PCR, expanded in LB broth with kanamycin, and purified using Qiagen Maxiprep kits (#12163). All constructs were verified by Sanger sequencing (Source Bioscience). Expression and puromycin resistance were validated in HEK293T cells prior to generation of stable SH-SY5Y cell lines.

### Generation of Stable Cell Lines

Stable SH-SY5Y cell lines were generated by plasmid transfection followed by antibiotic selection. Cells were seeded in 6-well plates and transfected the following day using 25 kDa linear polyethyleneimine (PEI; Alfa Aesar, #43896) according to standard protocols. Twenty-four hours post-transfection, cells were trypsinized and replated into 10 cm culture dishes to allow expansion prior to selection. Antibiotic selection was initiated 48 hours after transfection using the minimal concentration previously determined to eliminate non-transfected cells (typically 2 µg/mL puromycin [InvivoGen, ant-pr-1] or 100–200 µg/mL Geneticin [Gibco, 10131-035]). Selection-resistant colonies emerged within 2–6 weeks, depending on construct and antibiotic used. Monoclonal lines were established by isolating individual colonies using sterile filter paper discs soaked in trypsin, which were placed over selected colonies and transferred to individual wells for expansion. Polyclonal lines were generated by pooling multiple antibiotic-resistant colonies. For rapid generation of polyclonal puromycin-resistant SH-SY5Y lines, cells were alternatively seeded directly into 10 cm dishes and transfected at approximately 50% confluence. Forty-eight hours post-transfection, cells were subjected to cyclic puromycin selection consisting of 2 days of antibiotic treatment followed by 2 days of recovery in antibiotic-free medium. This selection cycle was repeated for approximately 2 weeks before transitioning to continuous puromycin maintenance.

### Stimulation of Cells with Light

Opsin-expressing cells were illuminated using a programmable LED array (NeoPixel Shield, Adafruit) controlled by an Arduino Uno. Blue (468.5 ± 1.5 nm), green (523.5 ± 1.5 nm), or red (622.5 ± 2.5 nm) LEDs were used as indicated. Culture plates were positioned 3 cm above the arrays using a custom 3D-printed gantry. Irradiance at the cell plane was empirically measured as 1.2 mW/cm^2^ (blue), 1.0 mW/cm^2^ (green), and 0.95 mW/cm^2^ (red). The Arduino program executed an initial 20 s delay after power-up, then continuously delivered the specified pulse parameters until power was removed.

For CoChR-RCaMP time-course experiments, blue and green LEDs were co-activated at maximum intensity at 8 Hz. Each flash comprised 20 ms of simultaneous blue + green illumination followed by 5 ms of green-only illumination. Flashes were delivered in 10 s trains (80 flashes) separated by 10 s inter-train intervals and repeated continuously for the stimulation period. Illumination was performed in a light-tight incubator maintained at 37 °C. Plates were housed in optically-isolated compartments to prevent cross-illumination, and each compartment was equipped with a continuously running ∼10 cm fan to minimize local heating. For long-term stimulation, culture medium was replaced with air-buffered medium (DMEM without NaHCO3 or phenol red supplemented with 10% FBS, 1% GlutaMAX, 1% non-essential amino acids, 1% penicillin-streptomycin, 0.2% MycoZap Plus, 5 g/L glucose, and 40 mM MOPS, pH 7.4), plates were sealed, and cells were maintained in darkness for 36 h prior to light exposure.

### Stimulation of Cells with Neurotransmitter Cocktail

To compare optogenetic excitation with receptor-mediated activation, cells were stimulated with a neurotransmitter cocktail modeled on wake-associated paradigms ^12^. A 100× stock solution was prepared in sterile water containing 1 mM carbachol, 100 μM NMDA, 100 μM AMPA, 100 μM kainic acid, 100 μM ibotenic acid, 100 μM serotonin, 100 μM histamine, 100 μM noradrenaline, 100 μM dopamine, and 1 μM orexin (all from Sigma unless otherwise specified). Aliquots were stored at −20°C. The stock was diluted 1:100 directly into culture medium to achieve working concentrations (1×), and cells were returned to the incubator for 4 h at 37°C. Following treatment, cells were lysed in TRI Reagent (Zymo Research, R2050) and total RNA was purified for downstream expression analysis.

### Quantitative Real Time PCR (qPCR)

Total RNA was isolated from cells using TRI Reagent (Zymo Research, R2050) followed by purification with the Zymo RNA extraction kit (R2052) or Zymo 96-well RNA kit (R2054) according to the manufacturer’s instructions, including on-column DNase digestion. RNA concentration and purity were assessed using a NanoDrop 1000 spectrophotometer (Thermo Scientific). For cDNA synthesis, 1 µg of total RNA was reverse transcribed using the High Capacity cDNA Reverse Transcription Kit (Applied Biosystems, 4368814) in a 20 µL reaction volume following the manufacturer’s protocol. After reverse transcription, cDNA was diluted to 100 µL with nuclease-free water, and 2 µL was used per qPCR reaction. Quantitative PCR was performed using the Universal Probe Library (UPL; Roche) system. Gene-specific primers were designed using the Roche UPL Assay Design Center, and appropriate UPL probes were selected according to manufacturer guidelines. Primer sequences and probe numbers are provided in Supplementary Table S1. Reactions were carried out using standard cycling conditions recommended for the UPL platform. Gene expression levels were quantified using the ΔΔCt method. Target gene expression was normalized to housekeeping genes within the same sample (GAPDH for SH-SY5Y cells; TBP for mouse-derived samples). Relative expression values were calculated as 2^−ΔΔCt^.

### Live-Cell Microscopy of RCaMP Expressing Cells

Live SH-SY5Y cells expressing RCaMP-containing constructs were imaged using a Leica SP8 confocal microscope maintained at 37 °C with 5% CO₂ conditions. Cells were imaged in air-buffered medium (see above) at 200 ms intervals. RCaMP fluorescence was excited using the White Light Laser (WLL) at 570 nm, and emission was collected between 580-620 nm using a hybrid detector. Pharmacological agents and salts were applied by addition of 2-20 µL of concentrated stock solutions directly to the imaging chamber. Optogenetic stimulation during imaging was delivered using the FRAP module. While continuously monitoring RCaMP fluorescence with low-intensity WLL excitation (570 nm), cells were exposed to high-intensity blue light pulses (400 ms) using the 488 nm and 476 nm argon laser lines to activate opsins. Frames acquired during blue light exposure were excluded from downstream analysis. Fluorescence traces were normalized using Leica software (maximum intensity scaled to 1, with linear scaling of remaining frames) and exported for analysis in R. The decay half-life of CoChR-mediated calcium transients was calculated using R scripts that identified individual peaks and determined the time required for signal amplitude to decrease to 50% and 25% of peak values.

### RNA-Seq Library Preparation and Sequencing

Total RNA was extracted from cultured cells using TRIzol reagent following two washes with PBS and purified using the Direct-zol RNA MiniPrep Kit (Zymo Research) according to the manufacturer’s instructions. RNA integrity was assessed using the QIAxcel system (Qiagen). RNA-seq libraries were prepared from 1 µg of total RNA using the KAPA RiboErase HyperPrep Kit (Roche) following the manufacturer’s protocol. Ribosomal RNA was depleted using complementary DNA oligonucleotides and RNase H-mediated degradation of RNA-DNA hybrids, followed by DNase treatment to remove residual DNA probes. rRNA-depleted RNA was fragmented at 94 °C for 6 minutes in 1× KAPA Fragment, Prime, and Elute Buffer to generate ∼200 bp fragments. Fragmented RNA was converted to double-stranded cDNA, followed by end repair, A-tailing, and adaptor ligation. Libraries were amplified by PCR using minimal amplification cycles (3-5 cycles) to reduce amplification bias. Following each enzymatic step, clean-up was performed using KAPA Pure Beads (Roche) according to the manufacturer’s instructions. Libraries were quantified and quality assessed prior to sequencing. Sequencing libraries were processed on Illumina HiSeq 4000 platforms to generate 100 bp paired-end reads.

### RNA-Seq Data Processing and Statistical Analysis

#### Read Alignment and Gene Quantification

Raw FASTQ reads were quality-filtered using Trimmomatic, applying sliding-window trimming (Phred <15 across 4 bases) and removing reads shorter than 36 nt. Reads were aligned to the human reference genome GRCh38 (Ensembl release 108) using STAR v2.7.10a with default parameters and --quantMode GeneCounts. Transcript assembly and quantification were performed using StringTie (v2.1.x) in reference-guided mode (-e flag) with the Ensembl 108 GTF annotation. Gene-level expression values were summarized as FPKM and TPM. For downstream analyses, gene-level FPKM matrices were used. Any transcript with a zero FPKM value at any time point was excluded from further analysis to eliminate genes expressed at very low levels. Gene-level expression values were transformed as log_2_(FPKM + 1). To isolate stimulation-dependent transcriptional responses, differential expression at each time point was calculated as Δlog₂(FPKM + 1) = light-stimulated (Light) – constant darkness (Control). Because inference focused on within-gene, within-timepoint Light–Control contrasts, conclusions relied on relative differences rather than absolute expression scaling.

#### Time-resolved differential expression and persistence

Time-resolved differential effects were assessed using a two-way ANOVA model applied independently to each gene: as log_2_(FPKM + 1) ∼ (time * condition), where timepoint (9 levels) and condition (Control vs Light) were fixed effects. The interaction term tested whether the effect of light stimulation varied across time. Post-hoc pairwise comparisons were performed using Tukey’s Honest Significant Difference (HSD) test to estimate Light–Control contrasts within each timepoint. P-values were adjusted using the Benjamini-Hochberg false discovery rate (FDR) procedure. For each gene, we computed: (i) The number of timepoints with significant Light–Control differences (“persistence”). (ii) The direction of each significant effect (Up if Light > Control; Down if Light < Control). Genes were binned by persistence (1, 2, or ≥3 significant timepoints). Directionality classes were defined as: Up-only: significant Up at ≥1 timepoint and never Down; Down-only: significant Down at ≥1 timepoint and never Up; Bidirectional: significant Up at ≥1 timepoint and Down at ≥1 timepoint across the series. The Bidirectional category captures direction switching and prevents double-counting that would occur when Up and Down persistence tables are merged. Genes with more than one zero value in either condition were excluded prior to modeling.

#### Circadian rhythm analysis

To evaluate rhythmic structure in control and light-stimulated conditions, we applied three complementary circadian detection methods to gene-wise expression time series (0-48 h, time points every 6 h, n=3 biological replicates per time point). RAIN was performed using the “independent” method to accommodate replicate samples per time point, testing for 24-h periodicity. P-values were adjusted using the Benjamini-Hochberg (BH) procedure. MetaCycle (meta2d) was used to integrate JTK_Cycle analyses with period constrained to 24 h. Integration statistics and method-specific parameters (phase, amplitude) were retained for downstream comparison. Harmonic regression was implemented by fitting cosine-sine models with a fixed 24-h period and comparing against a null (intercept-only) model using likelihood ratio tests. Amplitude and phase were derived from fitted coefficients, and BH correction was applied to yield corrected p-values. Genes passing method-specific adjusted thresholds were considered rhythmic. Overlap among methods was assessed descriptively; no integration score was used for classification.

#### Waveform-based temporal classification

To classify excitation-response kinetics, we analyzed gene-wise light-stimulated (BLUE) – constant darkness condition (CONT) trajectories within the 0–36 h window. For each gene, replicate-level expression was transformed as log_2_(FPKM+1), and Δ trajectories were computed per sample pair as Δ = BLUE − CONT at each timepoint. Gene-level mean and SEM of Δ were then computed across replicates at each timepoint. For waveform calling, per-gene Δ mean trajectories were smoothed across time using LOESS (degree 2; span = 0.8), yielding a smoothed series (Δ_s). From Δ_s, we extracted the maximum and minimum values (max_d, min_d) and their times (t_max, t_min), the fraction of timepoints below zero (frac_neg), the fraction of timepoints above a sustained threshold (frac_high), and the number of turning points (interior extrema) based on sign changes in first differences. Turning-point detection was made robust to small plateaus by forward-filling zero slopes; genes were required to have ≤3 turning points to be considered biphasic.

Waveform classes were assigned using explicit amplitude and shape gates applied to Δ_s: Suppression required a robust negative trajectory: min_d ≤ −0.25, max_d < 0.28, and ≥60% of timepoints below zero (frac_neg ≥ 0.60). This constraint prevents single-point dips from dominating classification. Biphasic (up→down) required a zero-crossing with both lobes exceeding thresholds (max_d ≥ 0.28 and min_d ≤ −0.25), correct ordering (t_max < t_min) with ≥6 h separation, a minimum negative-to-positive lobe ratio (|min_d|/max_d ≥ 0.45), limited turning points (≤3), and a rebound constraint such that the post-trough rebound did not exceed 45% of the positive lobe amplitude. Biphasic (down→up) required a zero-crossing with both lobes exceeding thresholds, correct ordering (t_min < t_max) with ≥6 h separation, and ≤3 turning points. Sustained induction required max_d ≥ 0.28 and persistence above a high fraction of the peak, defined as ≥50% of timepoints with Δ_s ≥ 0.65×max_d. Peak-timed induction classes were defined by the timing of the positive maximum among genes meeting the induction threshold (max_d ≥ 0.28): early peak if t_max < 24 h, mid peak if 24 ≤ t_max < 36 h, and late peak if t_max ≥ 36 h. Genes meeting the induction amplitude threshold but failing shape constraints were labeled Induction (weak shape), whereas genes failing amplitude/shape gates were labeled Other/weak. The Other/weak bin was treated as residual and excluded from downstream functional and regime-level interpretation. All waveform calls were performed on the smoothed Δ trajectories within the 0–36 h window, using fixed parameters across genes.

#### Functional and regulatory enrichment

Gene Ontology Biological Process (GO BP) enrichment was performed for each waveform class using hypergeometric testing (clusterProfiler), with all classified genes as the universe and Benjamini-Hochberg correction applied (mapped ENSEMBL to ENTREZ identifier). Transcription factor regulatory architecture was assessed using DoRothEA regulons (confidence levels A-C) after mapping ENSEMBL identifiers to gene SYMBOL. For each waveform class, overlap with TF target sets was evaluated using Fisher’s exact test. Enrichment effect sizes were calculated as log_2_(enrichment ratio) with gates of overlap a ≥ 3 and regulon size ≥ 10.

#### Cross-model generalization

Human SH-SY5Y waveform-classified genes were mapped to mouse orthologs using strict 1:1 Ensembl orthology, retaining only uniquely mapped gene pairs. For Hinard *et al.* ^12^, response genes (cortex and primary neuronal cultures) were intersected with ortholog-mapped genes, and per-waveform enrichment was tested using Fisher’s exact test with FDR correction. For the Diessler *et al.* ^19^, BXD panel, gene-wise sleep deprivation effects were computed as Δlog_2_CPM (SD–NSD) at ZT6. Genes were grouped by SH-SY5Y waveform class, and class-level summaries included distributional density plots, mean ± SEM SD effects, directional bias (fraction Δlog_2_CPM > 0; binomial test vs 0.5 with FDR correction), and tissue-paired effect sizes quantified using Cliff’s delta relative to the Other/weak class. For phenotype association analysis, strain-level gene scores were computed per waveform class and regressed against heritability-supported phenotype values (lm(value ∼ gene_score)). Regression coefficients (β) were z-scored within phenotype across waveform classes, averaged within phenotype categories (LMA, EEG, State), and rescaled for visualization.

#### Hallmark program scoring and temporal kernels

To summarize coordinated biological processes, MSigDB Hallmark gene sets were scored at each time point by computing gene-wise Z-scores across the time series and calculating the mean Z-score across genes within each set. Program-level temporal dynamics were modeled using kernel decomposition. For each Hallmark program, temporal response kernels were derived to capture recurrent activation motifs across lag time. Kernel polarity was defined as the integral of positive minus negative weights, and kernel centroid was computed as the lag-weighted mean of absolute kernel weights.

#### Hidden Markov modeling of regime structure

To infer higher-order transcriptional organization, we applied a Gaussian-emission hidden Markov model (HMM) to the matrix of Hallmark program activities across lag time. Models with varying state numbers were evaluated, and a three-state solution was selected based on stability and interpretability. Posterior state probabilities were estimated for each time point, yielding a probabilistic segmentation of the transcriptional response into discrete latent regimes. Programs were anchored to latent states using posterior-weighted contrasts (Cohen’s d), computed by weighting program activity by state posterior probability and comparing in-state versus out-of-state activity. Association strength was defined as |Cohen’s d|. State alignment was further quantified as the Pearson correlation between program activity and state posterior probability. Transition proximity was defined as 1 − max(posterior), providing a measure of uncertainty near regime boundaries.

#### Regime decomposition into waveform modules

To link regime-level organization back to gene-level temporal structure, state-specific program activity was decomposed into contributions from waveform-defined gene modules. Only genes assigned to structured waveform classes (excluding Other/weak) were included. For each Hallmark and waveform class: Gene-wise Z-scores were averaged within class. Contributions were weighted by class-specific gene counts. Count-weighting was used to reconstruct total program activity; qualitative state signatures were preserved under unweighted averaging. Weighted class contributions were verified to reconstruct global Hallmark activity. This decomposition quantifies how regime transitions arise from structured reweighting of temporally defined gene modules.

#### Statistical environment

All analyses were conducted in R (version 4.4.0) using tidyverse (version 2.0.0), limma (version 3.62.2), clusterProfiler (version 4.14.6), org.Hs.eg.db (version 3.20.0), msigdbr (version 25.1.1), dorothea (version 1.18.0), VennDiagram (version 1.8.2), ggplot2 (version 4.0.2), rain (version 1.40.0), MetaCycle (version 1.2.1), IMTest (version 1.0.0) and custom scripts. Hidden Markov modeling was implemented using Gaussian-emission models with posterior probability estimation. Multiple testing correction was performed using the Benjamini-Hochberg procedure unless otherwise specified.

## Results

### ChR2-mediated optogenetic stimulation elicits wake-associated transcription in human neuronal cells

We first asked whether controlled optogenetic depolarization is sufficient to induce activity-dependent transcriptional programs in human neuronal cells that resemble molecular signatures observed during wakefulness and sleep deprivation *in vivo*. SH-SY5Y cells were stably transfected with channelrhodopsin-2 (ChR2) and exposed to patterned blue light stimulation (Figure 1A; Supplementary Figure 1A).

**Figure 1.**
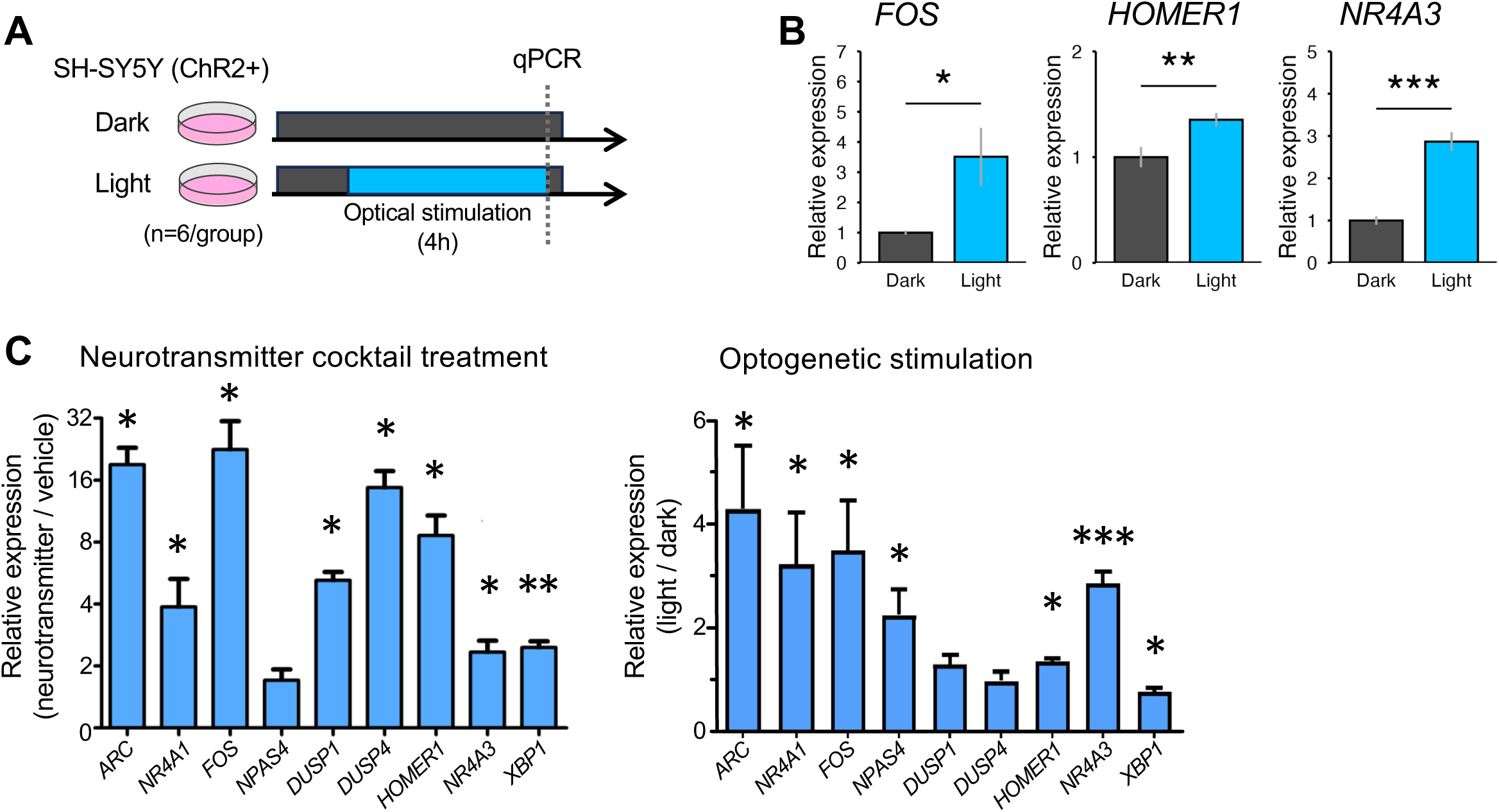
Optogenetic excitation of SH-SY5Y cells recapitulates sleep deprivation–associated gene induction. (A) Experimental schematic. SH-SY5Y cells stably expressing channelrhodopsin (ChR2) were subjected to patterned optical stimulation (Light) or maintained in darkness (Dark), followed by qPCR analysis. (B) Light stimulation significantly increased expression of canonical activity-dependent genes (*FOS*, *HOMER1*, *NR4A3*) relative to dark controls. (C) Comparison of gene induction following neurotransmitter treatment versus optogenetic stimulation. ChR2-mediated excitation robustly induced multiple sleep deprivation–associated immediate-early genes. Data are shown as mean ± SEM (n = 6 per group). Statistical significance: \**P* < 0.05, \*\**P* < 0.01, \*\*\**P* < 0.001.

Quantitative reverse transcriptase real-time PCR (qPCR) revealed significant induction of canonical activity-dependent genes, including *FOS*, *HOMER1*, and *NR4A3*, following light stimulation relative to dark-maintained controls (Figure 1B). These genes are well-established markers of neuronal activation and are induced in mouse cortex during sleep deprivation ^20^. The concordant direction of induction in SH-SY5Y cells indicates that optogenetic depolarization alone is sufficient to recapitulate core activity-dependent transcriptional responses observed during wakefulness *in vivo*. Among the tested genes, *FOS* exhibited the strongest induction, consistent with its role as an immediate-early gene and canonical readout of neuronal excitation.

Although the amplitude of induction was lower than typically observed with neurotransmitter cocktail treatment, light stimulation still increased target gene expression (Figure 1C), and the response was robust, reproducible, and strictly input-dependent. These data establish optogenetic stimulation of SH-SY5Y cells as an input-driven system capable of recapitulating core features of wake-associated transcription in isolation from systemic influences.

### Optimization of excitation parameters using high-sensitivity opsins

To enhance transcriptional output under prolonged illumination, we compared higher-sensitivity opsins: Chronos and CoChR, which exhibit greater photocurrent efficiency than ChR2 ^21, 22^. Both supported light-dependent *FOS* induction, but CoChR achieved robust activation at substantially lower stimulation frequencies (Supplementary Figure 2A,B). CoChR was therefore selected for subsequent experiments.

Calcium imaging (Supplementary Figure 2C) confirmed that CoChR stimulation engages physiologically relevant excitability pathways. SH-SY5Y cells exhibited no spontaneous calcium transients under baseline conditions but responded robustly to KCl depolarization (Supplementary Figure 2D) and excitatory neurotransmitters (Supplementary Figure 2E,F). Optogenetic stimulation evoked rapid, reversible intracellular calcium elevations (Figure 2A) whose amplitude scaled with light intensity (Figure 2B) and extracellular calcium concentration (Supplementary Figure 2G). At 8 Hz, three hours of stimulation induced transient *FOS* expression that decayed following light cessation, whereas six to nine hours produced sustained elevation (Figure 2C). Thus, transcriptional output scaled with stimulation duration, demonstrating that excitation history is encoded at the transcriptional level rather than reflecting only instantaneous depolarization. These data demonstrate that CoChR enables precise, reversible, and dose-controlled excitation in this system.

**Figure 2.**
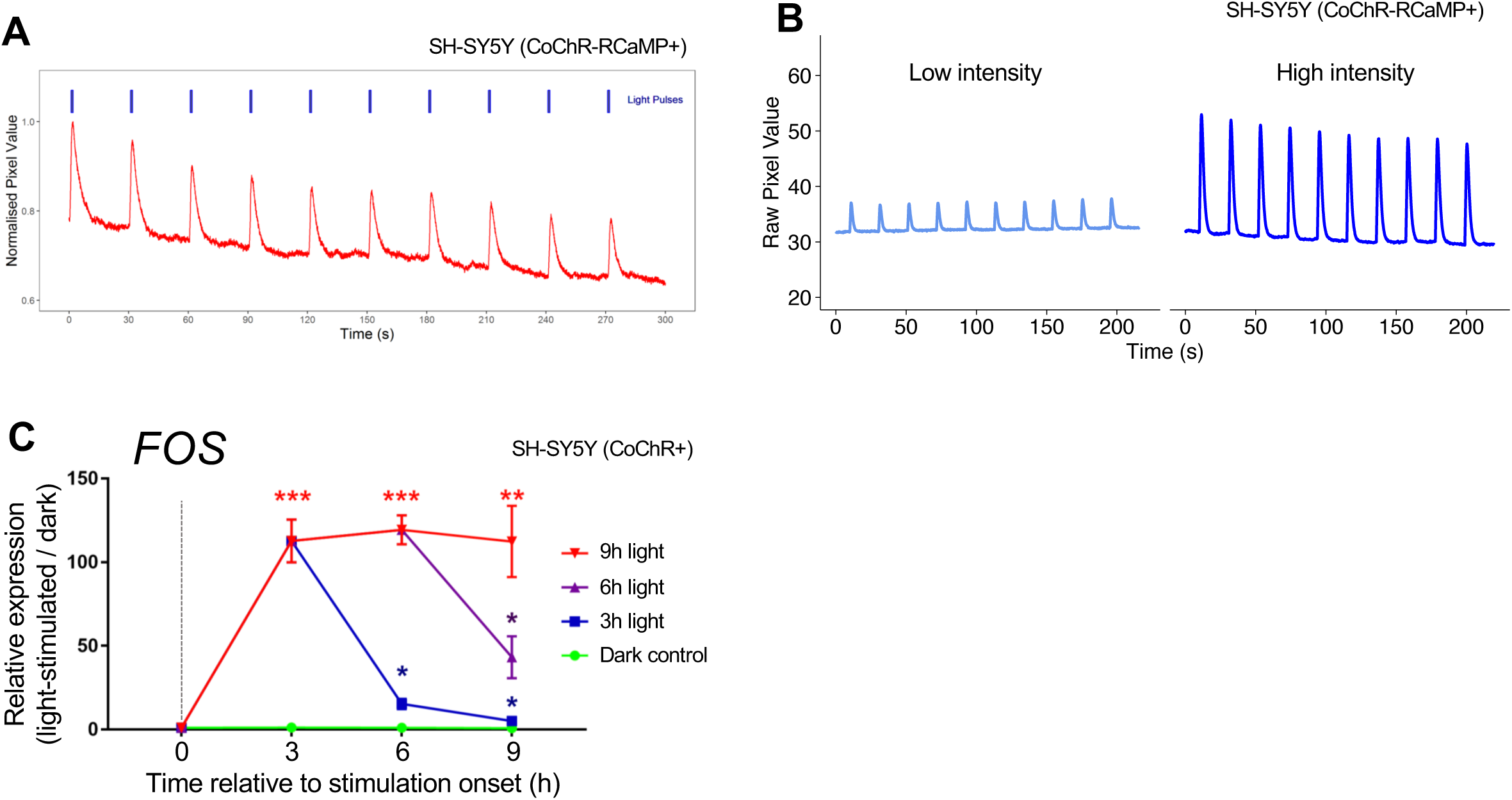
CoChR enables input-dependent calcium influx and sustained activity-induced gene expression in SH-SY5Y cells. (A) Patterned optical stimulation of CoChR-RCaMP–expressing SH-SY5Y cells induces rapid, repeatable intracellular calcium transients. (B) Light intensity–dependent calcium responses. Increasing light power produces larger-amplitude calcium transients, demonstrating graded input–output control of depolarization. Low intensity: 10% 488nm laser power; high intensity: (50% 476nm + 50% 488nm laser power). (C) Duration-dependent induction of the activity-responsive gene. SH-SY5Y cells expressing CoChR were stimulated at 8 Hz with alternating 470 nm and 570 nm light (total intensity 2.2 mW/cm²) for 3, 6, or 9 hours. *FOS* expression was quantified by qPCR (n = 4) relative to matched dark-treated controls. Short stimulation induces transient *FOS* elevation, whereas prolonged stimulation sustains high-level expression. Recovery time points are connected to the final illumination time point for clarity. Statistical significance is indicated relative to dark controls. \**P* < 0.05, \*\**P* < 0.01, \*\*\**P* < 0.001.

### Controlled excitation induces widespread and dynamic transcriptomic remodeling

Having established a robust excitation paradigm, we performed an RNA-seq time course to determine how sustained optogenetic activity reshapes global gene expression. SH-SY5Y cultures were synchronized, exposed to 12 hours of patterned light stimulation, and sampled every six hours over 48 hours alongside dark-maintained controls (Figure 3A).

**Figure 3.**
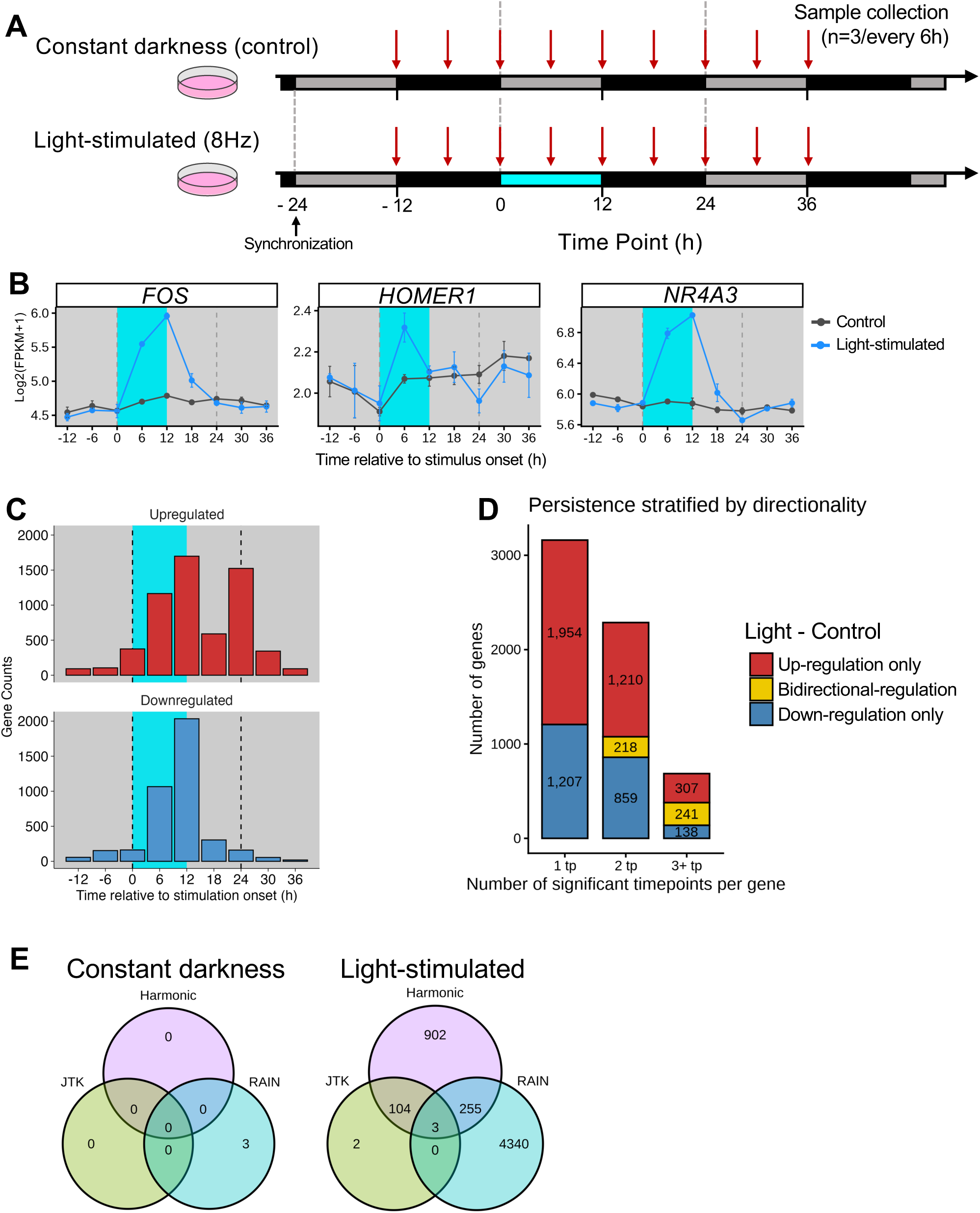
Sustained excitation drives temporally structured and predominantly transient transcriptional remodeling. (A) Experimental design. SH-SY5Y cells stably expressing CoChR were synchronized by serum shock and maintained in constant darkness (control) or exposed to patterned 8 Hz light stimulation for 12 h. Samples were collected every 6 h over a 48 h time course (n = 3 per time point). The shaded cyan region indicates the stimulation interval. (B) Representative activity-responsive genes illustrating temporal dynamics (log₂[FPKM + 1]). *FOS*, *HOMER1*, and *NR4A3* show robust induction during light exposure and return toward baseline after stimulation offset. (C) Distribution of peak induction timing for significantly upregulated and downregulated genes relative to stimulation onset. Most genes peak during or shortly after the stimulation window, indicating temporally structured activation. (D) Persistence of regulation stratified by directionality. Genes are grouped by the number of significant time points per gene (1, 2, or ≥3). The majority of regulated genes exhibit transient, single-timepoint responses, with fewer genes showing sustained or bidirectional regulation. (E) Circadian rhythmicity analysis under constant darkness and light-stimulated conditions using JTK_CYCLE, RAIN, and harmonic regression. Circadian detectors yield limited and inconsistent calls, consistent with structured but non-periodic excitation-driven dynamics rather than de novo circadian oscillations.

Canonical activity-dependent genes previously validated by qPCR in ChR2 cells behaved similarly in CoChR-expressing cells. *FOS*, *HOMER1*, and *NR4A3* were robustly induced during light exposure and returned towards baseline after illumination ceased (Figure 3B), confirming reproducibility across opsins. Across the 48-hour time course, 6,134 genes exhibited significant Light–Control differences at one or more sampling points (two-way ANOVA; Tukey HSD FDR < 0.05; Figure 3C). Most responses were transient, with 3,161 genes significant at a single timepoint, whereas 686 genes displayed persistent change across three or more intervals (Figure 3D; Supplementary File S1).

Stratifying these responses by directionality showed that the majority were unidirectional (Up-only or Down-only), but a subset exhibited bidirectional regulation, switching sign across the time course. Specifically, bidirectional genes were detected across persistence bins, indicating that some excitation-responsive transcripts undergo rebound or overshoot dynamics rather than a monotonic induction or suppression. Together, these results show that sustained excitation induces broad but predominantly transient transcriptional remodeling, with only a minority of genes displaying persistent change.

Light-stimulated cells exhibited temporally structured transcriptional dynamics. To determine if light-stimulation might lead to 24-hour patterning of profiles, we used three circadian detection methods (RAIN, JTK_Cycle, harmonic regression) to quantify this. Overlap among the methods was limited (Figure 3E), consistent with their differing abilities to detect sinusoidal (JTK_Cycle, harmonic regression) vs non-sinusoidal waveforms (RAIN) ^23–26^. Because excitation was delivered as a single prolonged stimulus rather than a repeating cycle, these patterns are unlikely to reflect canonical circadian oscillation, but rather represent structured but non-periodic excitation-driven responses, given that we were unable to detect rhythms in control (unstimulated) cells (Figure 3E).

### Waveform-based classification partitions excitation responses into coherent kinetic modules

Although useful, differential expression analysis only identifies which genes change, but not how those changes unfold over time. Similarly, circadian algorithms are optimized for detecting repeating periodic patterns and are not designed to resolve single-pulse, history-dependent kinetics. The coexistence of transient induction, delayed responses, and non-oscillatory dynamics suggested that excitation organizes transcription into distinct kinetic response programs.

To resolve this temporal architecture, we next applied waveform-based classification to systematically define the shape and timing of excitation-driven transcriptional responses. We classified genes according to the shape of their excitation-induced transcriptional trajectories across stimulation (0-12 h) and recovery (12-36 h), using smoothed Δlog_2_(FPKM+1) profiles (light-stimulated – constant darkness).

Waveform classification revealed a limited set of recurrent kinetic patterns, including early peak, mid peak, late peak, sustained induction, biphasic (up→down), biphasic (down→up), and suppression (Figure 4A,B; Supplementary File S1). Genes not meeting amplitude or shape criteria were assigned to an “Other/weak” residual category. This class was not interpreted as a discrete biological module but was retained as a reference group in cross-model and sleep-phenotype enrichment analyses to quantify relative effect sizes of structured waveform classes. Representative genes closely tracked their class-average trajectories (Figure 4C), confirming that assignments captured genuine kinetic structure.

**Figure 4.**
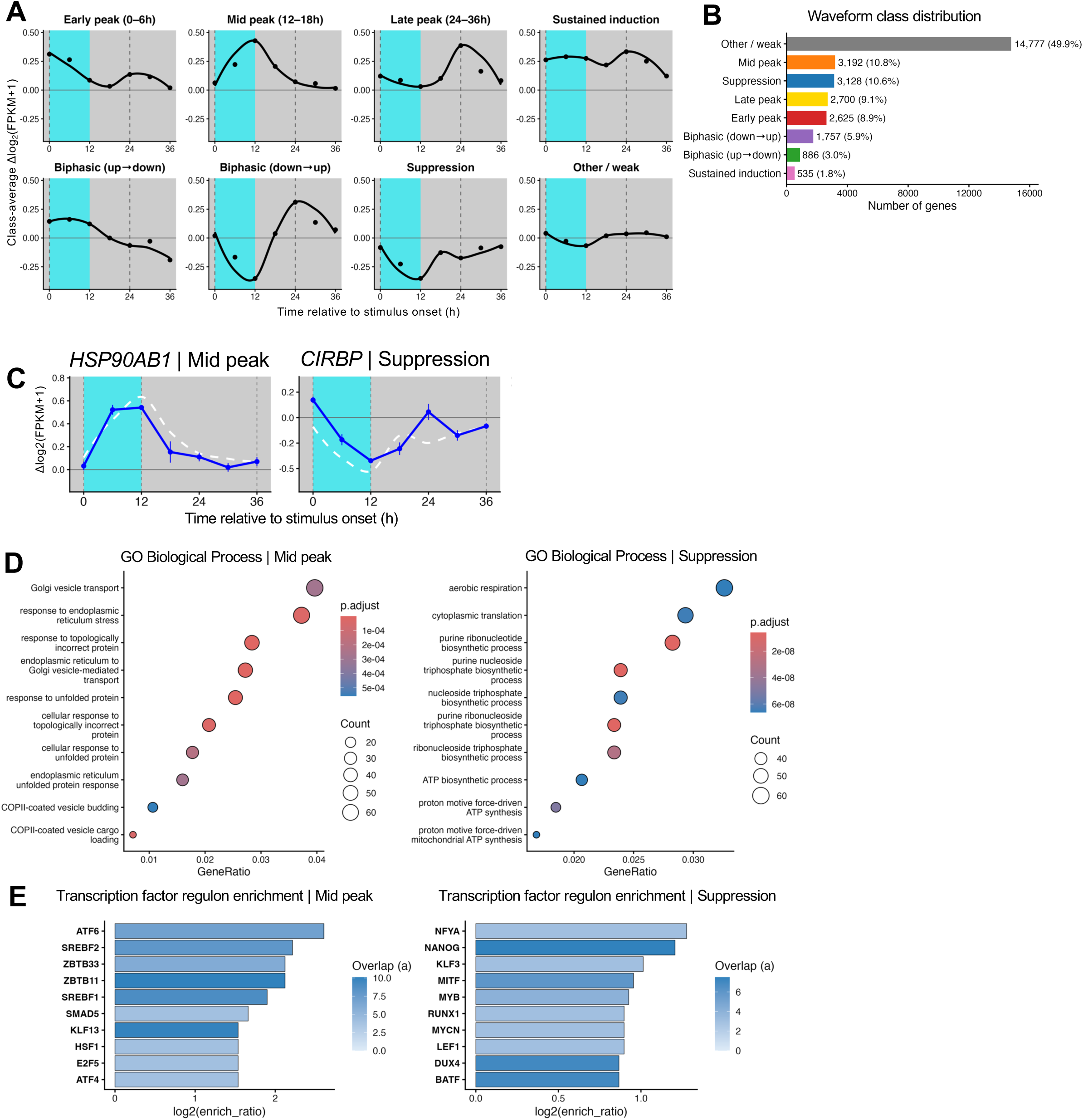
Excitation organizes post-stimulation transcription into discrete kinetic waveform modules with distinct functional and regulatory signatures. (A) Class-average Light–Control trajectories used for waveform assignment, computed from replicate-level expression as Δlog₂(FPKM+1) = Light − Control at each timepoint, then summarized across genes within each class (points = mean across genes). Classification was performed on the 0–36 h window using smoothed per-gene Δ trajectories (LOESS; span = 0.8). The cyan band denotes the light-stimulation interval, and dashed lines mark key boundaries used in the plotting frame. (B) Distribution of genes across waveform classes in the 0–36 h analysis window. Genes failing amplitude/shape gates were assigned to “Other/weak”, a residual bin for low-amplitude or irregular trajectories (not interpreted as a biological module in downstream analyses). (C) Representative genes illustrating distinct waveform behaviors (Mid peak and Suppression). Blue traces show mean Δlog₂(FPKM+1) with SEM; the dashed curve denotes the class-average smoothed trajectory for reference. (D) Gene Ontology (Biological Process) enrichment for example modules (Mid peak and Suppression), showing coherent functional specialization across kinetic classes (dot size = overlap/count; color = adjusted p-value; x-axis = GeneRatio). (E) DoRothEA transcription factor (TF) regulon enrichment highlighting distinct upstream regulatory signatures per module (examples shown for Mid peak and Suppression). TFs are ranked by effect size (log₂ enrichment ratio), with gates matching the analysis (minimum overlap a ≥ 3 and regulon size ≥ 10).

Functional enrichment analysis demonstrated that waveform classes corresponded to coherent biological programs (Supplementary File S1). For example, the Mid-peak class was enriched for protein trafficking and endoplasmic reticulum stress responses, consistent with delayed adaptive remodeling. In contrast, the Suppression class was enriched for biosynthetic and metabolic processes, consistent with compensatory downregulation following prolonged excitation (Figure 4D).

Transcription factor regulon enrichment further revealed distinct upstream regulatory signatures across classes (Figure 4E; Supplementary File S1). The Mid-peak module was associated with cell stress-adaptive regulators such as *ATF6*, *ATF4*, *HSF1*, and *SREBF* factors, whereas the Suppression module was linked to biosynthetic regulators including *NFYA* and *MYCN*. Thus, waveform classes represent coordinated kinetic programs defined by shared timing, function, and regulatory architecture. Waveform classification therefore resolves excitation-driven transcription into a limited set of temporally coherent modules that integrate kinetic shape, functional specialization, and regulatory architecture.

### Independent cross-model validation demonstrates conserved waveform programs

To test whether waveform-defined temporal modules reflect biologically meaningful programs rather than dataset-specific clustering, we evaluated each class against independent mouse sleep-wake transcriptomic datasets.

First, we mapped human SH-SY5Y waveform-classified genes to strict 1:1 mouse orthologs and tested their enrichment in cortical and primary neuronal culture sleep-deprivation (SD) responses from Hinard *et al.* ^12^. Because this dataset represents a single SD-versus-control contrast, we asked whether waveform identity predicts the direction of differential expression at the sampled time point. For each class, we compared the proportion of SD-up versus control-up genes using Fisher’s exact tests (2×2 contingency tables) and reported odds ratios with exact 95% confidence intervals (Figure 5A). Mid-peak and Suppression classes showed the strongest directional bias. This is expected: a static sleep-deprivation contrast captures a snapshot at one moment in time, and modules whose expression is maximally displaced from baseline at that moment will dominate the signal. Modules that peak earlier or later will be underrepresented, simply because they were sampled at the wrong phase. This sampling-phase dependence extends beyond these two classes, however. Directional structure was detectable across multiple module classes, indicating that static differential expression systematically reflects the underlying temporal identity of each transcriptional program.

**Figure 5.**
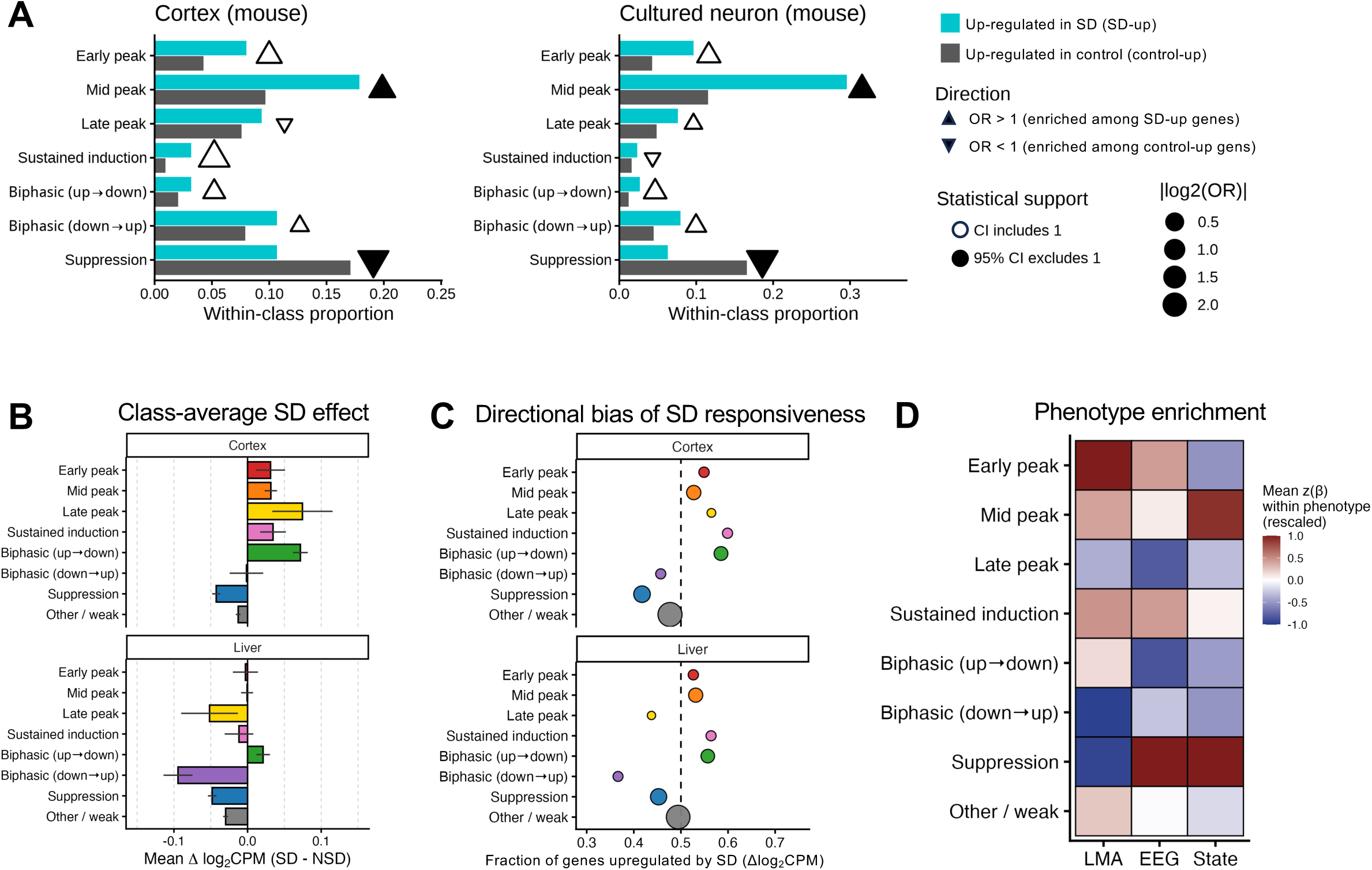
Temporal waveform classes generalize across species, experimental models, tissues, and sleep-related phenotypes. (A) Cross-species mapping of Hinard et al. mouse cortex and primary neuronal culture response genes onto human SH-SY5Y waveform classes using strict 1:1 orthology. Bars represent the proportion of stimulation-up (teal) and control-up (gray) genes assigned to each temporal waveform class in mouse cortex and cultured neurons. Triangles indicate odds ratios (OR) for enrichment of upregulated genes in sleep deprived group (SD-up) versus upregulated genes in control group (control-up) within each waveform (▲ OR > 1; ▼ OR < 1), with symbol size proportional to |log₂(OR)|. Filled symbols denote 95% confidence intervals excluding 1 (Fisher’s exact test). (B) Class-average SD effect (mean ± SEM across genes) for each waveform class in cortex and liver, highlighting distinct magnitude and direction of SD responsiveness by temporal class. (C) Directional bias of SD responsiveness by waveform class, shown as the fraction of genes with Δlog2CPM>0 (upregulated by SD) within each class (binomial test vs 0.5; multiple-testing corrected within tissue). (D) Temporal response class enrichment across heritability-supported phenotype categories in the BXD panel. For each phenotype, associations between phenotype values and waveform-class gene scores were fit across strains (lm(value ∼ gene_score)); β values were z-scored within phenotype across waveform classes and then averaged within phenotype category (LMA, EEG, State) and rescaled within category for visualization (mean z(β), rescaled).

We next tested whether temporal modules generalize beyond the tissue in which they were defined by analyzing a BXD sleep-deprivation dataset that profiled both cortex and liver (Diessler *et al.* ^19^). Liver provides a stringent test: if modules identified in cortex also organize transcriptional responses in a tissue with fundamentally different cellular composition and function, the underlying temporal programs likely reflect systemic regulatory logic rather than brain-specific circuitry. Gene-wise SD effects (Δlog_2_CPM, SD–NSD at ZT6) showed class-specific shifts in both tissues (Figure 5B). The ordering of these shifts mirrored intrinsic waveform phase: peak-type, sustained, and up→down modules shifted positively at ZT6, whereas down→up and Suppression modules shifted negatively (Figure 5C). Because effects were evaluated at a single timepoint, this ordering reflects where each module sits along its response trajectory at ZT6 rather than an absolute hierarchy of functional importance. Critically, although effect magnitudes were attenuated in liver relative to cortex, the directional ordering across classes was preserved, indicating that core temporal programs generalize beyond the central nervous system.

Finally, we asked whether waveform modules relate to genetically influenced organism-level traits. Using heritability-supported BXD phenotypes, we observed selective associations across locomotor activity (LMA), EEG, and behavioral state measures (Figure 5D). A functional ordering emerged. Early-peak modules preferentially associated with locomotor traits, consistent with rapid excitation-linked transcription coupling to phasic behavioral output. Sustained and mid-peak modules showed stronger associations with EEG and state phenotypes, aligning with prolonged network-level activity. In contrast, late-peak modules exhibited weaker and negative trait associations, consistent with roles in downstream adaptation rather than immediate behavioral output. Biphasic up→down modules were biased toward locomotor traits relative to EEG or state measures, suggesting transient activation followed by feedback repression influences dynamic behavioral transitions. Thus, waveform kinetics align with phenotype domain: rapid modules track phasic behavioral output, sustained/delayed modules align with network and state physiology, and late adaptive modules show minimal behavioral coupling.

Together, these analyses demonstrate that waveform classification of the cellular model transcriptional profiles captures conserved regulatory programs whose directional logic generalizes across species, tissues, and phenotypic scales. Temporal module identity therefore encodes biologically interpretable regulatory states rather than arbitrary clustering.

### Program-level dynamics reveal compositional organization of global Hallmark activity

To determine how sustained excitation restructures cellular-level function over time, we scored global activity of MSigDB Hallmark gene set programs across stimulation and recovery (Figure 6A; Supplementary File S1). Hallmark programs are curated signatures of well-defined biological states and processes, providing a standardized framework for summarizing complex transcriptomic changes into interpretable functional themes ^27^. Programs exhibited coordinated but non-uniform dynamics: growth- and cell-cycle–associated programs (E2F-targets, G2M-checkpoint, mitotic spindle) rose early during stimulation, inflammatory and stress pathways (TNFa signalings via NF-κB, unfolded protein response) peaked during the mid-phase, and metabolic and hypoxia-related programs became more prominent during recovery. Thus, these patterns revealed structured temporal organization at the systems level.

**Figure 6.**
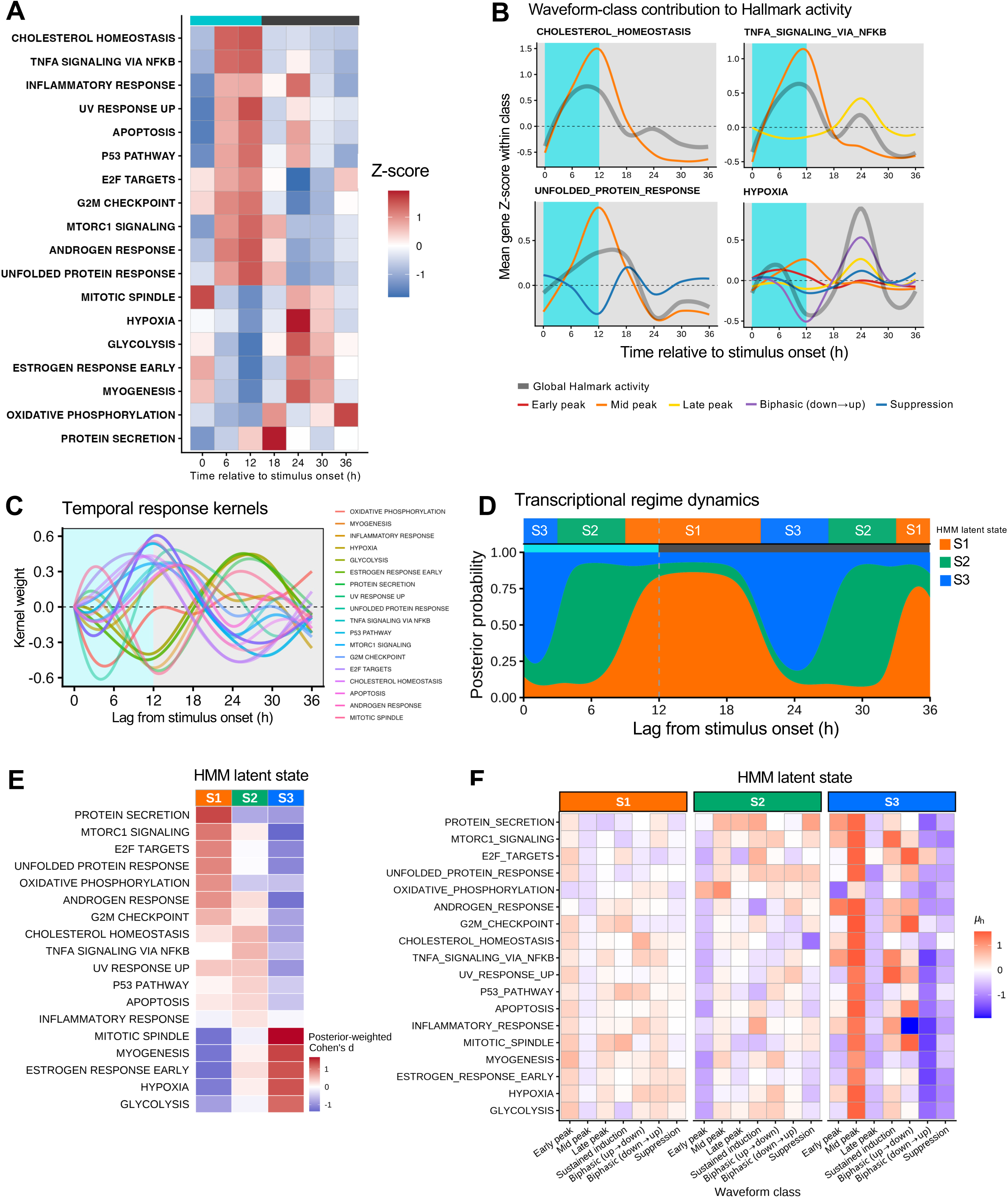
Excitation history organizes Hallmark programs into sequential global transcriptional states defined by distinct waveform-group compositions. (A) Global program dynamics. Z-scored Hallmark program activity (rows) across time relative to stimulation onset (columns), summarizing the global trajectory of coordinated biological programs during stimulation and recovery. (B) Relative waveform-class contributions to global Hallmark activity. Genes were first assigned to excitation-response waveform classes (Figure 4A). For each Hallmark gene set, we quantified the time-resolved contribution of each waveform class as the gene-count–weighted class mean Z-score (number of genes in class × mean gene-Z within class). These values represent the relative contribution of each kinetic gene subgroup to the global Hallmark trajectory. Waveform-class contributions were computed only for classes with sufficient representation and signal within each Hallmark set (≥15 genes and ≥50% of the maximal class amplitude range within that Hallmark). The black solid line denotes the reconstructed global Hallmark activity (mean gene-Z across all included genes), computed as the weighted mixture of class contributions. The shaded cyan region indicates the 0–12 h light-stimulation interval. Time is shown relative to stimulation onset. Thus, deviations in global Hallmark activity arise from dynamic redistribution of contributions across temporally distinct excitation-response gene classes rather than uniform regulation of pathway members. (C) Temporal response kernels learned for program-level dynamics, plotted as kernel weight versus lag (0–36 h). Shading indicates during-stimulation and post-stimulation intervals; dashed vertical lines mark stimulation onset and offset. (D) Sequential transcriptional state transitions. Hidden Markov Model posterior probabilities for three latent global transcriptional states (S1–S3) across lag time, showing ordered, directional state transitions induced by sustained excitation and maintained into recovery. The most likely state at each time point is indicated above. (E) State–program association map. Posterior-weighted Cohen’s d quantifying how strongly each Hallmark program is enriched or depleted within each state, defining state identity as a multivariate program configuration rather than single-pathway activation; negative values (blue) indicate relative depletion. (F) State composition by waveform groups. For each state (S1–S3), heatmaps show the waveform-group composition of Hallmark programs (mean standardized effect, μh), revealing how regime/state identity reflects the weighted contribution of underlying temporally structured gene groups. Together with (B), this links global state transitions to shifts in program composition, which in turn arise from redistribution of waveform-defined gene groups. Warmer colors indicate stronger positive contribution of a waveform class to a program in that state, whereas cooler colors indicate relative suppression.

Importantly, global program trajectories did not reflect uniform regulation of all genes within a pathway. Decomposition by excitation-response waveform class revealed that each Hallmark program is composed of temporally distinct gene subgroups whose weighted integration produces the observed global behavior (Figure 6B). For example, TNFa signaling via NF-κB showed an early rise driven predominantly by early-peak genes, whereas later dynamics reflected increasing contributions from late-peak modules. Similarly, unfolded protein response displayed strong mid-phase activation dominated by mid-peak genes, while suppression-class genes contributed opposing dynamics modulating the global activity. Pathway-level dynamics therefore arise from structured redistribution of temporally coherent gene modules within each program, rather than uniform regulation of all pathway members.

To formalize these patterns, we modeled program trajectories using temporal response kernels – smooth basis functions that capture canonical activation shapes such as transient peaks, sustained rises, or delayed responses (Figure 6C). Programs clustered into reproducible temporal motifs, indicating that excitation induces structured higher-order temporal organization rather than independent pathway fluctuations.

### Hidden Markov modeling identifies sequential global transcriptional regimes

Given the coordinated program dynamics, we next asked whether global activity could be summarized as transitions between recurring regulatory configurations. Hidden Markov modeling (HMM) of the Hallmark activity matrix identified three latent transcriptional regimes (S1–S3) that occurred sequentially across time (Figure 6D; Supplementary File S1). Transitions were probabilistic yet sequential (S1→S3→S2→S1), and did not simply reverse upon stimulation offset, indicating excitation-history–dependent reorganization rather than symmetric recovery.

Regime identity was defined by posterior-weighted program contrasts, illustrating how regimes are defined by coordinated clusters of functional programs (Figure 6E; Supplementary File S1). S1 was characterized by enrichment of protein secretion, mTORC1 signaling, E2F targets, unfolded protein response, and oxidative phosphorylation, with relative attenuation of mitotic spindle, myogenesis, hypoxia, and glycolysis. S3 showed an approximate inversion of this configuration, with mitotic and metabolic programs enriched and early growth-associated programs reduced. S2 exhibited a transitional configuration marked by comparatively elevated inflammatory and stress programs and reduced proliferative modules. Thus, each regime reflects a coordinated multivariate program configuration rather than isolated pathway activation.

### Global states emerge from structured reweighting of temporal gene modules

Because each Hallmark program comprises genes assigned to distinct excitation-response waveform classes (Figure 6B), we asked how regime identity relates to underlying gene-level timing (Figure 6F; Supplementary File S1). Partitioning state-specific program activity by waveform class revealed that global transcriptional states arise from coordinated reweighting of temporally defined gene modules embedded within each pathway.

Distinct redistribution patterns defined each state. In S1, early-peak modules contributed broadly across programs, consistent with acute excitation-driven activation, whereas mid-peak modules were relatively diminished. Late, biphasic, and suppression components were selectively retained within proliferative and metabolic programs (lower rows) but attenuated in stress-associated pathways (upper rows). In contrast, S3 was characterized by preferential enrichment of mid-peak modules and relative depletion of late-peak modules across proliferative and metabolic programs. Biphasic (down→up) and suppression modules were further reduced, particularly within proliferative and metabolic groups (lower rows), reinforcing a state-specific reconfiguration rather than uniform pathway regulation. S2 exhibited broadly reduced early-peak contributions and more diffuse waveform representation overall, consistent with a transitional configuration between early and late organizational states.

These results indicate that regime transitions reflect coordinated redistribution of temporally structured gene modules across functional programs. Sustained excitation does not uniformly activate or repress pathways; instead, it drives directional movement through regulatory states assembled from distinct temporal gene architectures, linking excitation-dependent gene kinetics to wake-associated systems-level reprogramming.

## Discussion

A major challenge in sleep-deprivation research is separating the direct molecular consequences of sustained neuronal activity from the systemic and behavioral changes that accompany prolonged wakefulness, including alterations in metabolism, temperature, hormonal signaling, and neuromodulatory tone ^28–33^. By imposing precisely controlled, reversible optogenetic excitation in a human neuronal-cell model, we isolate excitation history as the primary experimental variable. This reductionist framework reveals that sustained depolarization alone is sufficient to drive structured, directional transcriptional progression in the absence of glial, endocrine, or circuit-level inputs.

A central advance of this work is the demonstration that excitation-driven transcription is hierarchically organized. Gene-level responses segregate into temporally defined modules. These modules assemble within functional programs, and their shifting mixtures define discrete global transcriptional regimes. In this multilayered framework, cellular state reflects the weighted integration of temporally structured gene subsets rather than a uniform shift in pathway activity.

Notably, recovery did not retrace the induction trajectory in reverse. Instead, prior excitation biased subsequent transcriptional configurations, indicating that the system encodes cumulative activity history ^34, 35^. This asymmetry aligns with models of sleep pressure in which cellular demand reflects integrated prior wake rather than instantaneous firing rate alone ^36–38^. Excitation history may therefore leave a molecular imprint that shapes subsequent regulatory states.

The engagement of proteostatic and metabolic programs in this cell-autonomous context is particularly notable. Sleep deprivation *in vivo* has been associated with unfolded protein responses and metabolic strain ^39–42^, but these effects are often interpreted as secondary to systemic stress. Our findings demonstrate that sustained neuronal excitation alone is sufficient to initiate core components of these pathways. This is particularly relevant given growing evidence linking proteostasis failure to neurodegenerative disease ^6, 43–45^, and the association of sleep loss with impaired protein homeostasis and accumulation of misfolded proteins *in vivo* ^46–48^. Our data therefore supports a mechanistic link between cumulative neuronal activity burden and engagement of proteostatic stress networks. Moreover, modulation of heat-shock factor 1 (HSF1), a master regulator of the unfolded protein responses, alters sleep-wake phenotypes *in vivo* ^49^, supporting the view that proteostasis regulators function not merely as downstream markers of stress, but as active modulators of global state.

At the same time, the simplicity of the system defines its limits. The absence of glial interactions, inflammatory amplification, endocrine inputs, and circuit-level feedback means that the transcriptional configurations described here likely represent a foundational layer of regulation. In intact organisms, these intrinsic programs are expected to be further shaped by intercellular and systemic signals. Thus, the regimes identified in this study should be viewed as core excitation-driven states upon which additional biological complexity is superimposed.

More broadly, this work reframes activity-dependent transcription as progression through a structured state space defined by temporally organized gene modules. In this view, a transcriptomic snapshot encodes integrated excitation history, providing a mechanistic bridge between cellular activity dynamics and systems-level wake-associated reprogramming. Extending this framework to multicellular neuronal systems and organoids will allow excitation history to be experimentally imposed and state-space progression to be quantitatively mapped in more complex biological contexts.

## Acknowledgements

A.B.R. was funded by the Perelman School of Medicine, University of Pennsylvania, the EMBO Young Investigators Programme, and the Lister Institute of Preventative Medicine. This work was supported also by NIH DP1DK126167 and R01GM139211 (to A.B.R.).

## Author contributions

S.Y. performed the data analysis, synthesized findings, and wrote the manuscript.

W.G. designed the project, performed experiments and data analysis.

U.K.V. performed experiments and data analysis.

A.B.R. designed the project and analyzed results, secured funding, synthesized findings, and wrote the paper with contributions from the other authors.

All authors have reviewed the final manuscript and agree on its interpretation.

---

**Supplementary Figure 1.**
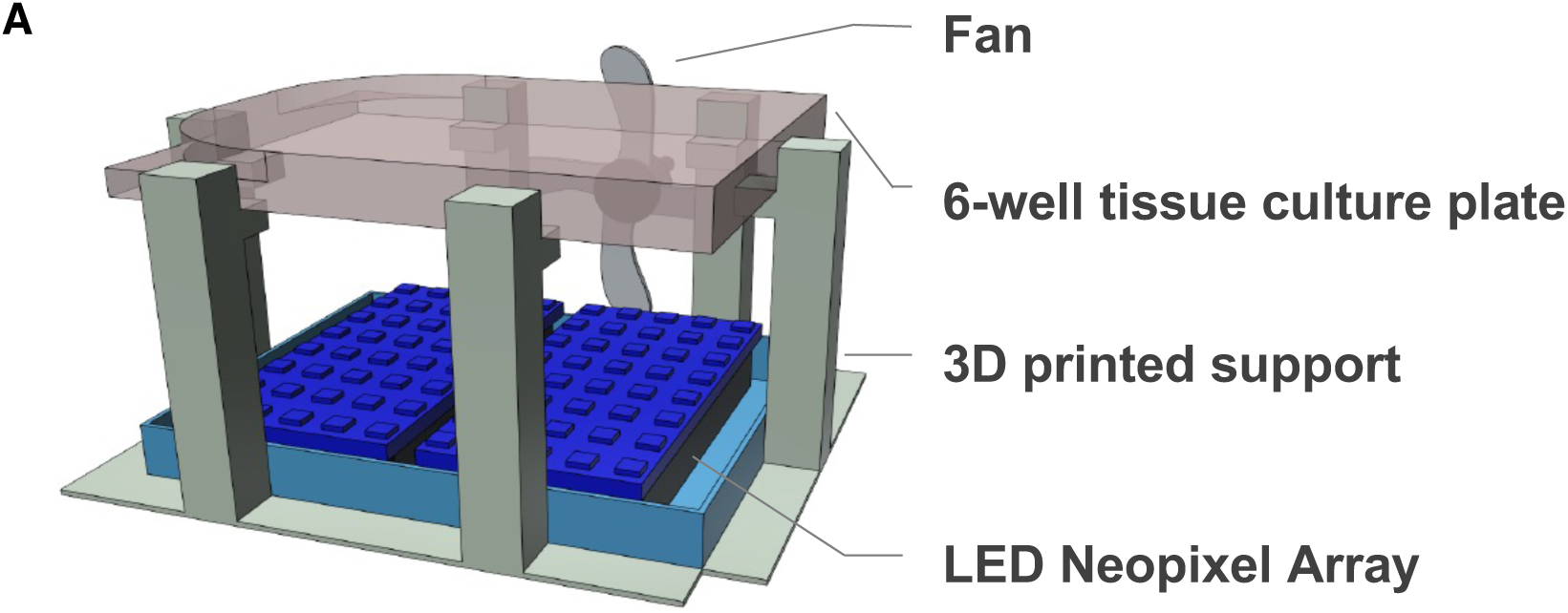
Custom LED stimulation platform. (A) Schematic of the custom LED-based illumination system. A 6-well culture plate was positioned ∼3 cm above an addressable RGB LED (NeoPixel) array mounted within a 3D-printed support. A cooling fan was installed above each plate to maintain temperature stability during illumination. Each array was controlled by an Arduino microcontroller and housed in individual light-tight compartments within a 37 °C incubator to prevent ambient light exposure. Power cycling every 10 minutes synchronized independent LED shields across arrays.

**Supplementary Figure 2.**
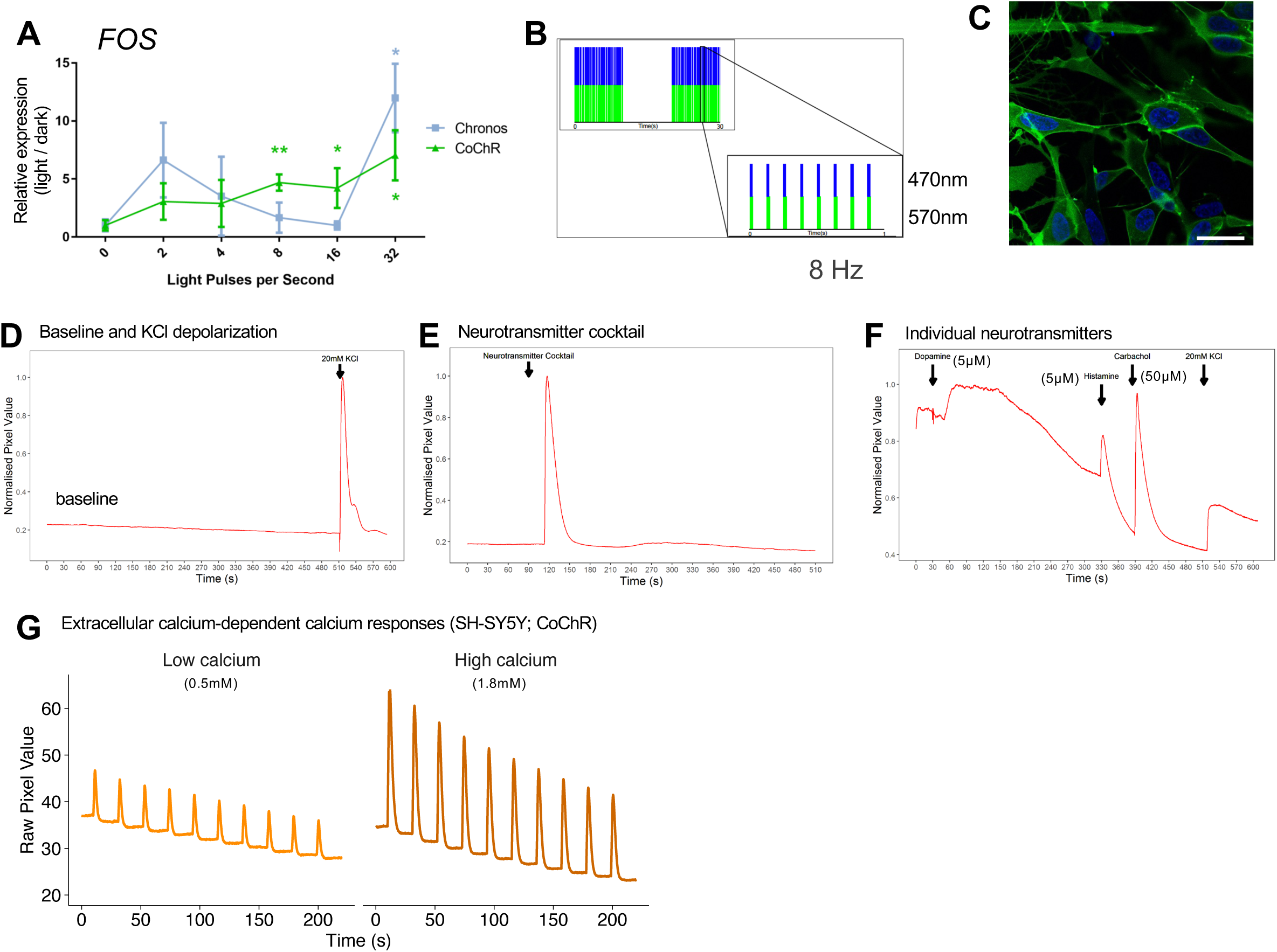
Validation of CoChR-mediated optogenetic stimulation and stimulus-dependent calcium signaling in SH-SY5Y cells. (A) Comparison of opsin performance. SH-SY5Y cells stably expressing Chronos or CoChR were stimulated with patterned blue/green light (6 h; total intensity 2.2 mW/cm²; 1 μM retinal). Fos expression was quantified by qPCR relative to matched dark controls (n = 3). CoChR exhibits stronger induction at lower stimulation frequencies compared with Chronos. (B) Schematic of patterned optical stimulation protocol (8 Hz; 470 nm and 570 nm light). (C) Representative fluorescence image of SH-SY5Y cells expressing CoChR-RCaMP. Scale bar: 20µm. (D) Baseline calcium activity in SH-SY5Y cells shows no spontaneous transients; depolarization with 20 mM KCl induces robust calcium elevation. (E) Application of an excitatory neurotransmitter cocktail induces transient intracellular calcium increases. (F) Individual neurotransmitters (dopamine, histamine, carbachol) elicit calcium responses, confirming functional neuromodulatory responsiveness. (G) Extracellular calcium–dependent calcium responses in CoChR-RCaMP–expressing SH-SY5Y cells. Lower extracellular calcium (0.5 mM) reduces calcium transient amplitude compared with physiological calcium (1.8 mM), consistent with calcium influx through light-gated channels. Statistical significance in (A) is indicated relative to matched dark-treated controls. *P < 0.05, **P < 0.01.

**Supplementary Table S1. Oligonucleotide sequences used in this study.**

Gibson_Primers: Primer sequences used for amplification and Gibson Assembly of opsin expression constructs, including ChR2(C128T), Chronos, CoChR, RCaMP1h, IRES-PuroR, and 2A-eNpHR2.0 fragments cloned into the modified pEGFP-N1 backbone. Primer sequences are listed 5′→3′ with corresponding target fragments and assembly context. qPCR_Primers: Gene-specific primer sequences used for quantitative real-time PCR (qPCR) assays. Primers were designed using the Roche Universal Probe Library (UPL) Assay Design Center; corresponding UPL probe numbers are indicated where applicable. Housekeeping genes (GAPDH for SH-SY5Y; TBP for mouse samples) are included for normalization.

**Supplementary File S1. RNA-seq temporal classification and regulatory analyses.**

Waveform_classification: Per-gene temporal classification based on smoothed Light–Control (Δlog₂(FPKM+1)) trajectories within the 0–36 h window (spanning stimulation onset through 24 h post-stimulation recovery). Features include amplitude metrics, turning-point counts, biphasic shape constraints, suppression robustness criteria, and final waveform class assignment.

GO_BP_enrich_wave: Gene Ontology Biological Process enrichment results for each waveform class using hypergeometric testing with Benjamini–Hochberg correction. Universe defined as all waveform-classified genes.

TF_enrich: Full transcription factor enrichment analysis per waveform class using DoRothEA regulons (confidence levels A–C). Enrichment statistics include Fisher’s exact test p-values, adjusted FDR, enrichment ratios, and effect sizes.

TF_enrich_top_effect_size: Subset of transcription factor enrichments ranked by effect size (log₂ enrichment ratio), highlighting dominant regulatory associations per waveform class.

HMM_posterior_probability: Posterior probabilities from Gaussian-emission hidden Markov modeling of Hallmark program activity across lag time. For each lag, probabilities of latent states (S1–S3) are provided and sum to 1.

HMM_Hallmark_anchor: State anchoring metrics for each Hallmark program. Includes posterior-weighted in-state versus out-of-state contrasts (Cohen’s d), correlation with state probability, and absolute anchor strength.

HMM_Hallmark_wave_decomposition: Decomposition of state-specific Hallmark activity into contributions from waveform-defined gene modules, quantifying how temporal gene classes are reweighted across latent regimes.

